# Programmable protein editing by split intein-mediated recombination

**DOI:** 10.64898/2026.01.22.700961

**Authors:** Masaharu Somiya, Takumi Yanase

**Author notes:** Correspondence to Masaharu Somiya.

## Abstract

Biological regulation has long relied on modifying genes, transcripts, or protein abundance, yet direct rewriting of protein sequence after translation remains largely inaccessible. Here, we introduce a programmable protein editing platform based on split intein-mediated recombination, enabling precise excision and replacement of defined segments within mature proteins in living cells. By covalently exchanging a target segment with a donor-encoded sequence supplied in trans, this approach functions as a protein-level analogue of recombination, operating independently of transcription or translation. Using ultrafast and orthogonal split inteins, we demonstrate efficient protein recombination in mammalian cells, enabling single-amino acid substitutions, domain replacement, functional switching, and reprogramming of subcellular localization. This work establishes post-translational protein recombination as a general strategy for sequence-level control of protein function, expanding the conceptual and practical scope of biological editing beyond nucleic acids.

## Introduction

Proteins are composed of modular domains and finely tuned sequence motifs that together determine their structural and functional properties. Although genome editing enables precise manipulation of DNA/RNA sequences^1–3^, no general technology currently exists for post-translationally rewriting protein sequences to repair, reprogram, or replace specific segments within polypeptides. A platform capable of editing proteins directly without modifying the genome would provide powerful opportunities for conditional control of protein function, rapid exploration of sequence-function relationships, and development of new therapeutic modalities.

Protein trans-splicing (PTS) mediated by split inteins offers a rare biochemical mechanism for nearly traceless covalent ligation of polypeptides. Natural split inteins such as Npu DnaE^4,5^ and gp41-1^6,7^ catalyze peptide bond formation between two extein fragments while excising themselves from the final product (**Figure 1A**). Existing PTS applications have leveraged this chemistry to assemble split proteins^8^, perform segmental isotopic labeling^9,10^, generate cyclic proteins^11^, induce protein alternative splicing ^12^, or conditionally activate engineered constructs^10,13^. Recent studies have shown that split inteins enable the insertion of a short segment into a loop region of target protein^14,15^, which resembles protein editing to some extent, and reconstruction of large functional dystrophin by trans splicing that effectively treats dystrophic mice^16^. However, the core capability of inteins, the ability to mediate precise covalent replacement, has not been fully exploited for editing proteins.

**Figure 1.**
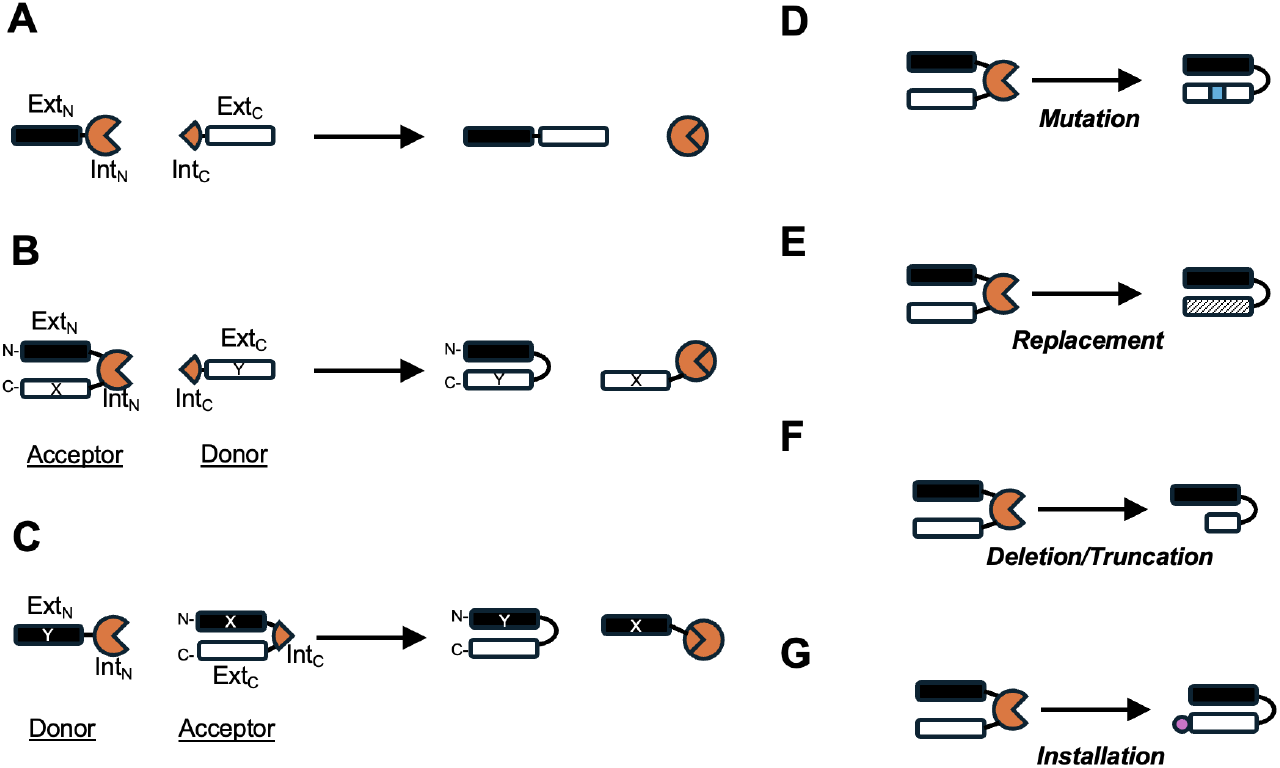
Scheme of split intein-mediated protein editing. (A) Canonical split intein-mediated trans-splicing. The N- and C-intein fragments (Int_N_ and Int_C_) are fused to the corresponding N- and C-exteins (Ext_N_ and Ext_C_), and trans-splicing ligates Ext_N_-Ext_C_ while excising the intein fragments as a separate product. (B) Domain replacement by placing Int_N_ in the acceptor. The acceptor contains Ext_N_ fused to Int_N_ and a target domain X; the donor carries Int_C_ fused to the replacement domain Y. Trans-splicing yields the edited product in which X is replaced by Y, and the excised byproduct contains the intein fragments together with domain X. (C) Alternative architecture in which Int_C_ is placed in the acceptor between domain X and Ext_C_, enabling replacement of an N-terminal domain X with donor-derived domain Y. (**D to G**) Mutation enables substitution of a single or a few amino acids (D). Replacement enables the exchange of a larger segment or structured domain (E). Deletion/Truncation enables complete excision of a target domain by using a donor lacking a replacement sequence, or partial deletion by providing a shortened fragment as the donor payload (F). Installation enables the addition of new elements ranging from small epitope tags to large structured domains (G).

A compelling conceptual opportunity is to use engineered inteins to excise a segment of a target protein (acceptor) and replace it with a designer sequence supplied in trans (donor). By placing intein insertion sites at permissive positions, this approach can produce a continuous edited polypeptide while releasing the original protein segment (**Figure 1B and 1C)**. Such a system would transform inteins from simple ligation modules into a general-purpose platform for protein sequence editing. Importantly, the donor sequence introduced during trans-splicing is not constrained to resemble the endogenous segment. Therefore, the same framework supports multiple scales of editing, enabling a continuum of applications, such as single-residue precision editing, replacement of entire structured domains, deletion of specific regions, or installation of short tags or large domains (**Figure 1D to 1G**). These editing modes dramatically extend the utility of intein chemistry beyond conventional applications and establish a biochemical analogue to CRISPR-style genome editing—but acting directly on mature proteins rather than on genetic material. Recent discoveries and engineering of split inteins, particularly gp41-1 and Npu derivatives^16,17^ with relaxed extein specificity and faster kinetics, now make these concepts practicable. These inteins can splice efficiently even when flanking residues deviate from canonical extein identities, enabling insertion into structured regions and compatibility with a wide range of editing payloads.

In this study, we demonstrate a generalizable post-translational protein editing platform based on split inteins. By pairing an acceptor protein harboring a native segment with a donor fragment carrying an engineered sequence, we achieve efficient trans-splicing that yields an edited protein in mammalian cells. Critically, we validate that this system supports editing at multiple scales, from single-residue replacement to large-scale sequence rewriting and complete domain substitution, highlighting its versatility and modularity. This work establishes the conceptual and technical foundation for a new class of biochemical technologies capable of post-translational protein sequence editing. Such capabilities expand the toolkit for synthetic biology, mechanistic protein analysis, new therapeutics, and modular redesign of protein architectures.

## Results

### Trans splicing-mediated recombination of split luciferase

To establish a quantitative reporter for split intein-mediated protein editing, we adopted the split NanoLuc luciferase (Nluc) system^18^. Nluc can be reconstituted from an inactive large fragment (LgBiT) and an 11-amino acid peptide (HiBiT) that binds LgBiT with subnanomolar affinity (K_D_ = 700 pM). As a stringent negative control, we used DrkBiT, which differs from HiBiT by a single amino acid substitution; DrkBiT retains binding to LgBiT but does not restore luciferase activity, providing a “binding-competent but catalytically dark” control^19-21^. We designed an intein-mediated recombination assay in which trans-splicing converts DrkBiT into a functional HiBiT sequence. Specifically, the acceptor protein comprised LgBiT fused to the N-terminal fragment of the GP41-1 split intein (Int_N_), followed by the inactive peptide DrkBiT (LgBiT-Int_N_-DrkBiT). The donor fragment consisted of the complementary C-terminal intein fragment fused to HiBiT (Int_C_-HiBiT). In this architecture, productive trans-splicing excises the DrkBiT-containing segment and installs the donor-encoded HiBiT sequence in its place, thereby generating an active LgBiT-HiBiT luciferase (**Figure 2A**).

**Figure 2.**
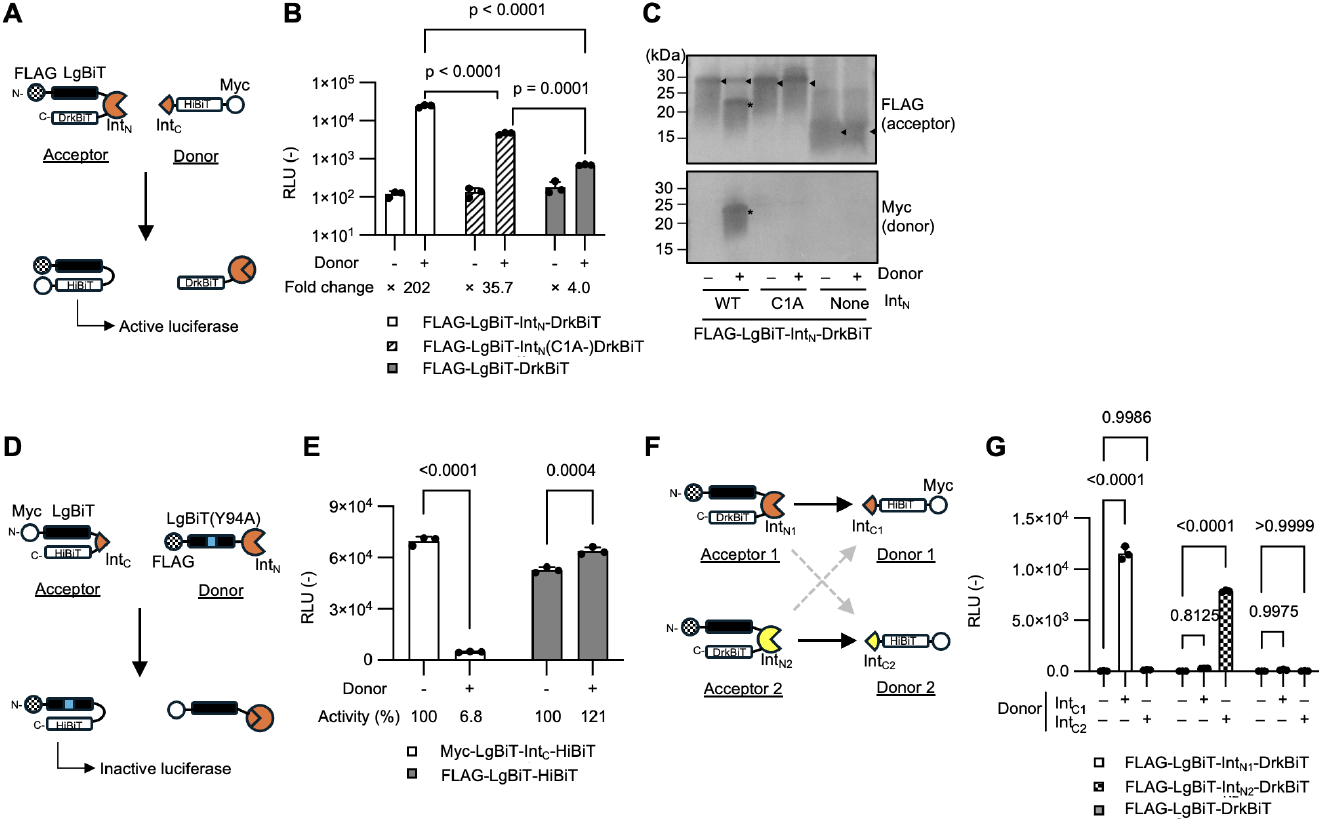
Split intein-mediated recombination of NanoBiT reporters. (A) Schematic of a NanoBiT activation design in which recombination between FLAG-LgBiT-Int_N_-DrkBiT (acceptor) and Int_C_-HiBiT-Myc (donor) yields an LgBiT/HiBiT-containing enzymatically active product. Sequences of HiBiT and DrkBiT are VSGW**<u>R</u>**LFKKIS and VSGW**<u>A</u>**LFKKIS, respectively. (B) Nluc activity (RLU) in transfected HEK293T cells measured for the recombination conditions in (A) using Int_N_(WT), Int_N_(C1A), and no-Int_N_/no-donor controls. Bars show mean ± SD (N=3). Fold-change (FC) relative to the no-donor condition is shown below the axis. Exact P values from two-way ANOVA with Tukey’s multiple-comparisons test are indicated. (C) Western blot analysis detecting acceptor (FLAG) and donor (Myc) species under the conditions in (B). (D) Schematic of a NanoBiT inactivation design in which recombination between Myc-LgBiT-Int_C_-HiBiT (acceptor) and FLAG-LgBiT(Y94A)-Int_N_ (donor) yields an inactive LgBiT(Y94A)/HiBiT configuration. (E) Nluc activity measured for the conditions in (D) (± donor). Bars show mean ± SD (N=3). Activity (%) compared to the no-donor condition is shown below the axis. Exact P values from two-way ANOVA with Tukey’s multiple-comparisons test are indicated. (F) Orthogonality test schematic: two acceptors carrying distinct Int_N_ fragments (IntN_1_ or IntN_2_) were paired with cognate donors carrying Int_C1_ or Int_C2_; only matched Int_N_/Int_C_ pairs are expected to splice and convert DrkBiT into HiBiT to activate Nluc. (G) Nluc activity for the orthogonality matrix in (F) in transfected HEK293T cells. Bars show mean ± SD (N=3). Exact P values from two-way ANOVA with Tukey’s multiple-comparisons test are indicated.

When expressed without donor in HEK293T cells, LgBiT-Int_N_-DrkBiT did not produce detectable Nluc activity, consistent with the requirement for both halves and for catalysis. In contrast, co-expression of the acceptor and donor induced a robust increase in luminescence (**Figure 2B**), indicating efficient sequence replacement sufficient to restore enzymatic activity. To confirm that the observed signal depended on intein catalysis rather than noncovalent complementation or fragment stabilization, we introduced a catalytic-inactivating point mutation in the acceptor intein (Cys1 to Ala; C1A), which has been shown to abolish GP41-1 splicing^6,13^. The C1A mutation greatly decreased luminescence rescue upon co-expression (**Figure 2B**), demonstrating that trans-splicing is required for recombination-mediated editing in this system. Notably, a substantial residual signal persisted with C1A, consistent with preserved Int_N_-Int_C_ binding that tethers donor HiBiT near acceptor LgBiT and permits weak proximity-driven complementation despite abolished splicing. As expected, when the acceptor protein lacks the Int_N_ portion, there was only a negligible luciferase activity detected. This indicated that spontaneous exchange between DrkBiT in the acceptor and HiBiT in the donor without split intein-mediated binding or recombination did not contribute to a substantial restoration of enzymatic activity (**Figure 2B**).

To directly verify covalent recombination, we performed immunoblotting of cell lysates using an epitope tag (Myc tag) placed on the donor fragment. Co-expression yielded a new band at the expected molecular weight for the edited product, consistent with installation of the donor-derived sequence and tag into the acceptor protein (**Figure 2C**). The edited band was absent in the Int_N_(C1A) and no-Int_N_ controls, further supporting a splicing-dependent mechanism.

We next asked whether the same strategy can replace a substantially larger protein segment, rather than a short peptide. We therefore implemented an N-terminal recombination configuration in which the donor fragment carries an N-terminal extein (Ext_N_) fused to Int_N_ (Ext_N_-Int_N_), while the acceptor protein contains Int_C_ positioned downstream of active LgBiT (LgBiT(WT)-Int_C_-Ext_C_). In this orientation, trans-splicing installs the donor-encoded N-terminal region (Ext_N_) of inactive LgBiT(Y94A) onto the acceptor, effectively replacing a large N-terminal segment in a single editing event and inactivating luciferase activity (**Figure 2D**). Co-expression of these components dramatically decreased Nluc activity by 93% compared to the control (**Figure 2E**), indicating that the edited product is almost completely inactivated after replacement of a large N-terminal domain. As before, the lack of Int_C_ in the acceptor prevented recombination-mediated inactivation (**Figure 2E**), confirming that editing depends on intein catalysis rather than simple fragment association.

Finally, to demonstrate modularity and assess orthogonality, we implemented the same NanoBiT editing architecture using an independent split intein pair (Ava_N_/Npu_C_^22^). Matched Int_N_ /Int_C_pairs supported robust Nluc activation, whereas mismatched combinations failed to generate a signal (**Figure 2F and G**), indicating minimal cross-reactivity between intein systems. These results support the feasibility of using multiple orthogonal split inteins to enable parallel or multiplexed protein editing within the same cellular context.

Collectively, these split luciferase experiments establish a quantitative, splicing-dependent platform demonstrating that split intein-mediated protein editing can encode mutation (DrkBiT to HiBiT), replacement (N-terminal swapping), and sequence installation (tagging) through donor design (**Figure 1**).

### Split intein-mediated editing of functional proteins for activity switching

To extend intein-mediated editing beyond Nluc complementation to a functional protein, we used mNeonGreen (mNG), a bright monomeric green-yellow fluorescent protein^23^. The mNG chromophore is derived from the tripeptide motif GYG at residues 58-60. We introduced an inactivating AAA substitution at this site (mNG(AAA)), which blocks chromophore maturation and eliminates fluorescence. We then designed an editing strategy in which trans-splicing replaces the mutated chromophore-encoding segment with a donor fragment restoring the native GYG sequence (**Figure 3A**). To enable this replacement, we inserted Int_C_ at position 146, a surface-exposed loop between the 7th and 8th β-strands of mNG. Upon co-expression of the acceptor and donor, robust fluorescence was recovered, whereas no signal was detected when the acceptor was expressed alone or when intein catalysis was disabled by the C1A mutation (**Figure 3B and 3C**).

**Figure 3.**
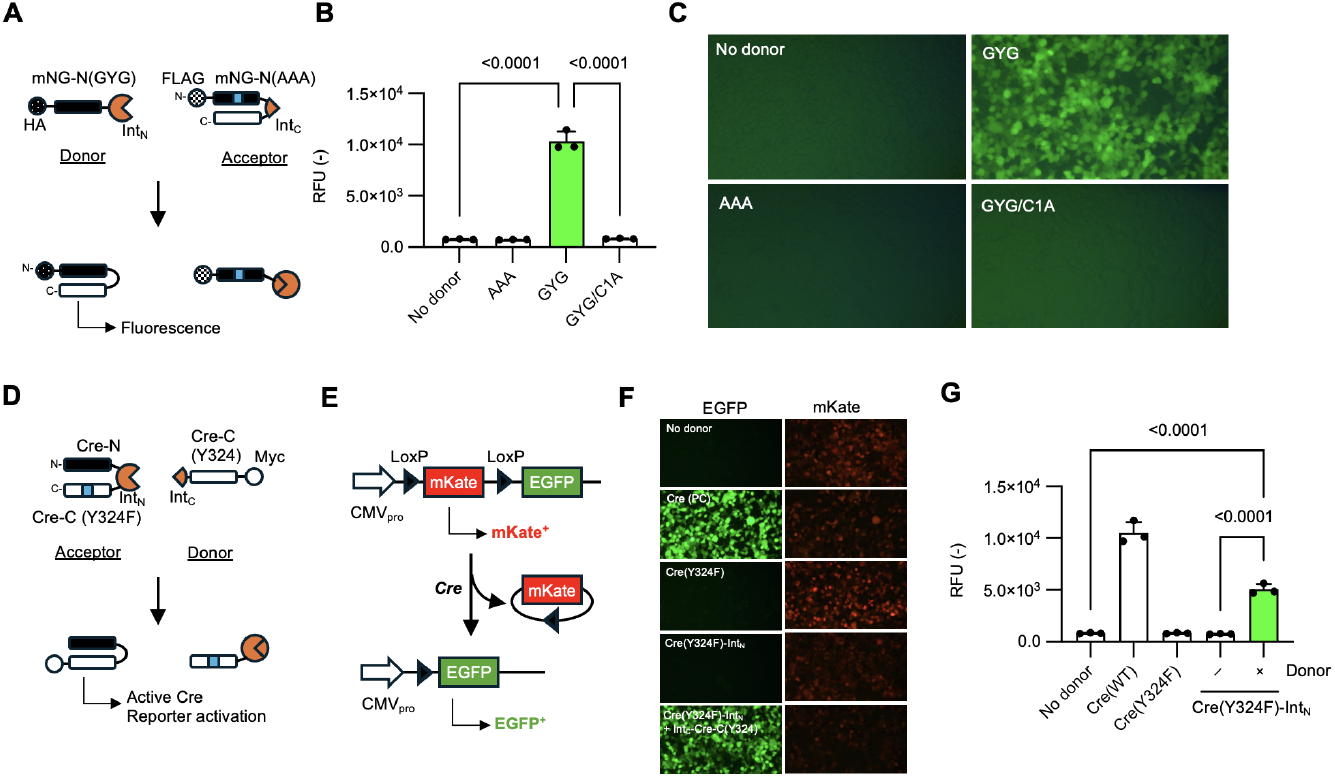
Functional switching of mNeonGreen and Cre recombinase by intein-mediated protein editing. (A) Schematic of the mNeonGreen (mNG) editing assay. A non-fluorescent acceptor construct (mNG-FL(AAA)-Int_C_) is combined with a donor fragment (mNG-N(GYG)-Int_N_) to generate a fluorescent mNG product. (B) Quantification of mNG fluorescence in HEK293T cells. The acceptor protein (FLAG-mNG-N(AAA)-Int_C_) was expressed alone (no donor) or co-expressed with donor fragments encoding mNG-N(AAA)-Int_N_, mNG-N(GYG)-Int_N_, or mNG-N(GYG)-Int_N_(C1A). Bars show mean ± SD (N = 3). Exact P values from ANOVA with Tukey’s multiple-comparisons test are indicated. (C) Representative fluorescence microscopy images of HEK293T cells corresponding to the conditions in (B). (D) Schematic of split Cre editing. The acceptor encodes Cre-N fused to Int_N_ and an inactive C-terminal fragment Cre-C(Y324F); the donor encodes Int_C_-Cre-C(Y324). (E) Schematic of the Cre reporter assay. Cre activity induces recombination of a loxP-flanked reporter cassette, resulting in a switch from mKate to EGFP expression. (F) Fluorescence images of the Cre reporter assay in HEK293T cells. EGFP and mKate channels are shown for cells expressing wild-type Cre (positive control), Cre(Y324F), the acceptor alone, or acceptor plus donor. (G) Quantification of Cre reporter activation based on EGFP fluorescence for the conditions in (F). Bars show mean ± SD (N = 3). Exact P values from ANOVA with Tukey’s multiple-comparisons test are indicated.

We next asked whether intein-mediated editing can switch the activity of a widely used enzyme in mammalian genetics, Cre recombinase^24,25^. Cre activity can be eliminated by well-characterized point mutations, including R173K and Y324F^26,27^. We constructed an inactive Cre(Y324F) acceptor and supplied a donor fragment encoding the corresponding wild-type sequence to restore Cre via trans-splicing (**Figure 3D**). To enable efficient fragment reconstitution, we inserted Int_N_ at the canonical split-Cre junction between residues 59 and 60 (Asn59/Asn60), a site previously shown to support robust complementation upon reassembly^28^. Cre function was evaluated using a loxP reporter in which Cre-mediated recombination switches expression from mKate to EGFP (**Figure 3E**) ^29^. Co-expression of acceptor and donor induced strong reporter activation, whereas acceptor alone failed to activate the reporter (**Figure 3F and 3G**). Together, these results show that split intein-mediated recombination can restore enzymatic activity in cells by rewriting defined residues within a functional protein.

### Post-translational protein editing rewires functional state and cellular behavior

To create a “turn-on” editing assay with a secreted readout, we fused LgBiT to Int_N_, the inhibitory peptide DrkBiT, and appended an ER retention signal (KDEL^30^) (LgBiT-Int_N_-DrkBiT-KDEL), yielding an inactive reporter confined to the ER. In this configuration, DrkBiT blocks productive luciferase complementation, while KDEL enforces ER retention. The donor was designed to drive GP41-1-mediated recombination that replaces DrkBiT with HiBiT to activate the reporter and excise the KDEL-containing C-terminus to permit secretion. In the combined mode, a single editing event simultaneously installs HiBiT and removes KDEL, generating an active, secretion-competent LgBiT-HiBiT complex (**Figure 4A**).

**Figure 4.**
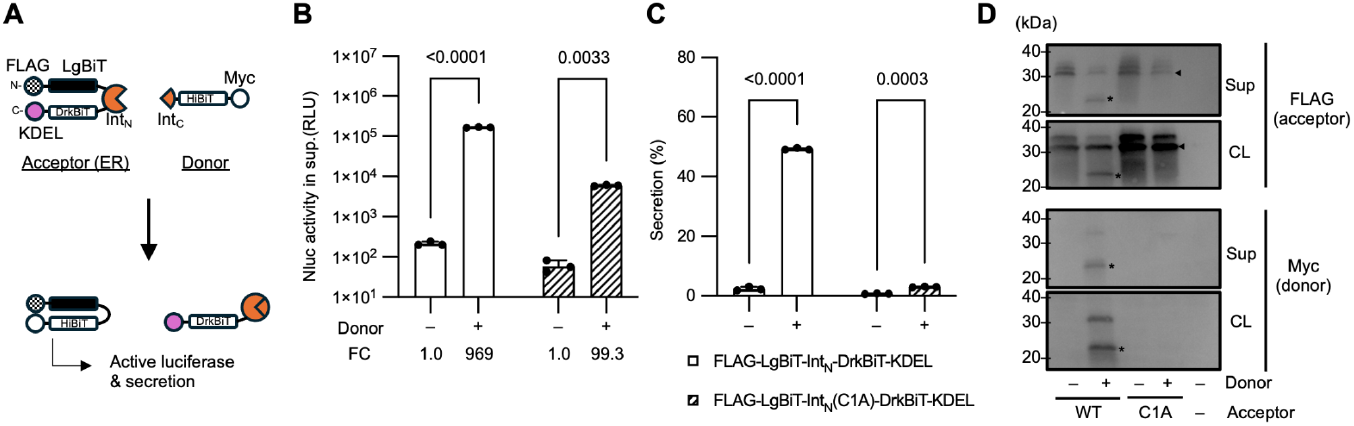
Secretion assay for intein-mediated protein editing of an ER-retained reporter. (A) Schematic of the secretion-on editing assay. ER-retained FLAG-LgBiT-Int_N_-DrkBiT-KDEL (acceptor) is edited by HiBiT-Int_C_-Myc (donor) via recombination to convert from DrkBiT to HiBiT and remove KDEL, generating active luciferase and secretion. (B) Nluc activity measured in culture supernatants for cells expressing the donor construct with either the wild-type acceptor (open bars) or a splicing-defective acceptor (Int_N_(C1A); hatched bars). Bars show mean ± SD (N=3). Fold-change (FC) relative to the no-donor condition is shown below the axis. Exact P values from two-way ANOVA with Tukey’s multiple-comparisons test are indicated. (C) Secretion efficiency quantified as the percentage of total reporter activity detected in the supernatant [(supernatant) / (supernatant + lysate) × 100] for the same conditions as in (B). Bars show mean ± SD (N=3). Exact P values from two-way ANOVA with Tukey’s multiple-comparisons test are indicated. (D) Western blot analysis of cell lysate (CL) and supernatant (Sup) fractions to verify intein-mediated product formation and donor-dependent changes in ER retention/secretion (probed via the indicated epitope tags).

In HEK293T cells, this combined editing induced robust luciferase activation and shifted the signal from cell-associated to supernatant (**Figure 4B**). Quantification of luminescence and western blotting of lysate and conditioned medium confirmed that an intact catalytic residue (Cys1 in Int_N_) was required (**Figure 4B to 4D**). Together, these data demonstrate that intein-mediated protein editing can concurrently reprogram protein activity and subcellular localization—in this case, converting an inactive, ER-tethered protein into an active, secreted form—using a single supplied donor.

### Donor-encoded rules enable multiplex rewriting of protein localization

To test whether a single donor can rewrite localization determinants across multiple targets in parallel, we constructed three acceptor reporters with distinct initial destinations: Myc-mNG-GP41-1-N-Fis1 (Fis1-derived targeting signal for mitochondrial outer membrane^31^), HA-mTagBFP2-GP41-1N-NES (nuclear export signal (NES) for cytosol localization^32^), and HiBiT-mScarlet-GP41-1-N-CAAX (CAAX motif for plasma membrane anchoring^33^). Each acceptor places GP41-1-N immediately upstream of a C-terminal targeting element (Fis1, NES, or CAAX), such that editing swaps the targeting element without altering the FP core (**Figure 5A**).

**Figure 5.**
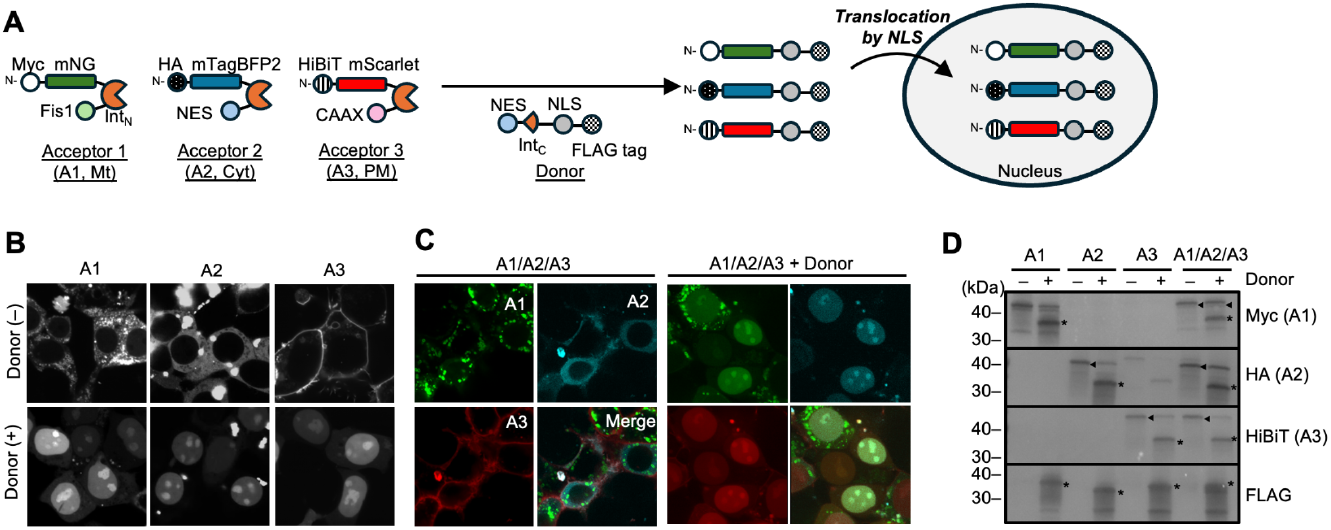
Subcellular relocalization by intein-mediated protein recombination. (A) Multiplex relocalization by a single donor. Three acceptor proteins bearing Int_N_ fragments are targeted to distinct compartments via built-in localization signals—mitochondria (Acceptor 1, A1), cytosol (Acceptor 2, A2), and plasma membrane (Acceptor 3, A3). A common donor (NES-Int_C_-NLS-FLAG) drives intein-mediated recombination that replaces the C-terminal localization module of each acceptor, thereby removing the original targeting signal and installing an NLS, resulting in convergent nuclear accumulation of all three fluorescent reporters. (B) Confocal images of HEK293T cells expressing each acceptor in the absence or presence of the donor, showing donor-dependent nuclear relocalization. (C) Co-expression of acceptors A1/A2/A3 with the donor illustrates simultaneous conversion of all three reporters to a common nuclear destination. (D)Western blot analysis confirming donor-dependent recombination product formation for each acceptor, detected via the indicated epitope tags (Myc, HA, HiBiT, and FLAG). Asterisks indicate edited products; triangles indicate acceptor proteins.

We used a single donor encoding NES-GP41-1C-NLS-FLAG, which is designed to trans-splice onto each acceptor and replace its original C-terminal signal with a shared SV40 NLS^34^. When expressed alone, each acceptor localized to its anticipated compartment: mitochondria, plasma membrane, or cytosol. Upon donor expression, all three reporters redistributed to the nucleus, indicating efficient rewriting of three distinct C-terminal targeting signals into a common nuclear destination (**Figure 5B and 5C**). Consistent with the expected trans-splicing products, western blotting showed that edited proteins retained their N-terminal epitope tags (Myc, HA, or HiBiT) while gaining the donor-encoded FLAG tag, confirming precise installation of the NLS-FLAG segment at the C terminus (**Figure 5D**). As anticipated for productive trans-splicing, the edited bands migrated at a lower apparent molecular weight than the corresponding acceptors, reflecting removal of the acceptor-encoded Int_N_ fragment and its original targeting signal and replacement with the NLS-FLAG segment. Simultaneous editing of all three reporters was also demonstrated under multiplexed conditions (**Figure 5C and 5D**). Together, these results demonstrate multiplex, donor-programmed relocalization using a single universal donor architecture.

## Discussion

Two recent papers^14,15^ highlighted the power of split inteins as “protein editing” tools, using trans-splicing to rewrite short internal segments of target proteins—primarily to install small cargos such as epitope tags, short peptides, or chemically functionalized elements, using two orthogonal pairs of split inteins. These studies established intein-mediated editing as a precise and efficient strategy for localized protein modification. By contrast, our work develops a more general editing architecture in which the donor sequence is freely designable, enabling edits across scales—from single-amino acid substitutions to domain-scale replacement, deletion, or truncation—and allowing multiple features to be encoded within a single editing event. This flexibility enables the combinatorial rewrites of protein sequences and states in a single step, expanding intein-mediated editing from localized installation to general-purpose protein rewriting.

These properties make intein-mediated editing a powerful addition to the synthetic biology toolbox^35,36^. Beyond basic research, a modular architecture also creates opportunities to couple editing to other regulatory inputs, such as small molecules, external light stimuli, or engineered sensing domains. Because the system is genetically encodable, delivery as DNA or mRNA could enable programmable, in situ switching of protein states in vivo. Moreover, unlike genome editing, this strategy does not require permanent alteration of the underlying gene sequence; thus, edited states can be transient and may avoid risks associated with irreversible genomic changes.

Conceptually, the reaction mechanism also supports an analogy to DNA recombination^37,38^: the excised acceptor segment and the introduced donor segment are designed to be complementary alternatives for the same position in the protein, such that the donor effectively “replaces” the original sequence while the removed fragment is released as a distinct product.

Despite these advantages, there are important limitations. The most fundamental constraint is that endogenous proteins cannot be edited directly unless the target protein is engineered to contain the split intein fragments at the desired positions; in other words, this platform currently requires prior genetic installation of intein components into the acceptor protein^14^. A second limitation is the possibility of residual intein-derived scars remaining in the edited product with some intein systems. Scarless inteins have been reported (e.g., variants such as Npu/Dna_E_ in prior work^16^), suggesting that this issue may be mitigated by intein choice and engineering. Relatedly, intein reactions can impose extein sequence requirements at the splice junctions. However, inteins with relaxed extein constraints have been described^16,17^, raising the possibility that junction requirements could be broadened and that fully scarless editing may become feasible across more sequence contexts. Finally, the fate of the excised fragments warrants consideration. The cleaved N- or C-terminal remnants released during editing could, depending on the application, interfere with the edited protein’s function^39^ (e.g., via residual binding, dominant-negative competition, or sequestration). In many cases, however, such fragments may be unstable in cells and subject to degradation over time^40^; nevertheless, this should be evaluated empirically for each target and application.

## Materials and Methods

### Biological materials and cell culture

Synthetic genes were obtained from Integrated DNA Technologies and GenScript. Synthetic genes were cloned using a type IIs restriction enzyme BsaI^41^, with mammalian expression vector pCMV or bacterial expression vector pET. For subcloning of genes, we used Gibson assembly^42^. Plasmids used in this study will be available through Addgene. HEK293T cells (obtained from ATCC) were cultured in Dulbecco’s Modified Eagle’s Medium supplemented with 10% fetal bovine serum at 37°C with 5% CO_2_.

### Cell-based assays for protein editing

HEK293T cells were transiently transfected with plasmid DNA using polyethyleneimine (PEI) as described previously^43^. Briefly, cells were seeded on day 1, transfected on day 2, and analyzed on day 3 or 4. For Nluc assays, cells were lysed 18–24 h post-transfection, incubated with furimazine, and luminescence was measured on a BioTek Synergy H1 plate reader (Agilent). For the reporter secretion assay, both culture supernatant and cell lysate were collected and subjected to the Nluc assay as described above. For immunoblotting, cells were lysed in RIPA buffer, clarified lysates were resolved by SDS-PAGE, and transferred to PVDF membranes. Membranes were probed with HRP-conjugated anti-DYKDDDDK antibody (FUJIFILM Wako, clone 1E6), anti-Myc antibody (MBL, clone My3), anti-HA antibody (MBL, clone TANA2), or anti-HiBiT antibody (Promega, clone 30E5) with an HRP-conjugated anti-mouse IgG secondary antibody (TCI) where applicable, and detected by chemiluminescence.

HEK293T cells were transiently transfected as described above and plated on glass-bottom dishes or multi-well slide chambers. At 24–48 h post-transfection, cells were imaged live. Fluorescence images were acquired using an epi-fluorescence microscope (SXJ-5822FLM, WRAYMER). Confocal images were acquired using a laser-scanning confocal microscope (FV-1000, Olympus) equipped with a 100× oil-immersion objective. Image processing and analysis were performed using Fiji/ImageJ. The fluorescence signal originating from transfected cells was measured on a BioTek Synergy H1 plate reader.

## Acknowledgments

We thank Toshihide Okajima and Yoh Wada at SANKEN, the University of Osaka, for helpful discussions. This work was supported by the Japan Society for the Promotion of Science KAKENHI (25H02269 to M.S.), Japan Science and Technology Agency PRESTO (JPMJPR24OB to M.S.), TERUMO LIFE SCIENCE FOUNDATION (24-III3009 to M.S.), Takeda Science Foundation (M.S.), and Astellas Foundation for Research on Metabolic Disorders (2024A2123 to M.S.). The authors acknowledge the use of ChatGPT (OpenAI) for assistance in improving language. All scientific content was generated, verified, and approved by the authors.

